# Unigene-based RNA-seq provides insights on drought stress responses in *Marsdenia tenacissima*

**DOI:** 10.1101/390963

**Authors:** Heng-Ling Meng, Wei Zhang, Guang-Hui Zhang, Jian-Jun Wang, Zhen-Gui Meng, Guang-Qiang Long, Sheng-Chao Yang

## Abstract

*Marsdenia tenacissima* is a well-known anti-cancer medicinal plant used in traditional Chinese medicine. Drought severely affects production and no information on its transcriptional responses to drought stress is available. In this study, cDNA libraries on control (CK), drought stress (T1), and re-watering (T2) treatments were constructed and HiSeq 2000 sequencing was performed using the Illumina platform. There were 43,129,228, 47,116,844, and 42,815,454 clean reads with Q20 values of 98.06, 98.04, and 97.88, respectively. A total of 8672, 6043, and 6537 differentially expressed genes (DEGs) were identified when CK vs. T1, CK vs. T2, and T1 vs. T2, respectively, were analyzed. In addition, 1039, 1016, and 980 transcription factors (TFs) were identified in CK, T1, and T2, respectively. Among them, 363, 267, and 299 TFs were identified as DEGs in CK vs. T1, CK vs. T2, and T1 vs. T2, respectively. These differentially expressed TFs mainly belonged to the bHLH, bZIP, C2H2, ERF, MYB, MYB-related, and NAC families. A comparative analysis of CK vs. T1 and T1 vs. T2 found that 1174 genes were up-regulated and 2344 were down-regulated under drought stress and this pattern was the opposite to that found after re-watering. Among the 1174 genes up-regulated by drought stress, 64 were homologous to known functional genes that directly protect plants against drought stress. Furthermore, 44 protein kinases and 38 TFs with opposite expression patterns under drought stress and re-watering were identified, which are possibly candidate regulators for drought stress resistance in *M. tenacissima*. Our study is the first to characterize the *M. tenacissima* transcriptome in response to drought stress, and will serve as a useful resource for future studies on the functions of candidate protein kinases and TFs involved in *M. tenacissima* drought stress resistance.

## Introduction

Drought is one of the most severe threats to crop production worldwide. It causes considerable yield losses and effects food security (Walter et al., 2011). Global warming means that drought will occur more frequently and will affect crop production more severely (Walter et al., 2011; Liu et al., 2015). Therefore, developing drought-tolerant crops is currently one of the main objectives of breeding programs. However, a deeper understanding of the molecular mechanisms underlying drought tolerance in crops is essential if new varieties with improved drought resistance are to be developed.

Over the last decade, the molecular mechanisms underlying plant drought tolerance have been widely investigated in different species using gene microarrays (Seki et al., 2002; Rabbani et al., 2003; Aprile et al., 2009; Luo et al., 2010; Le et al., 2012). As a result, thousands of genes have been identified that respond to drought stress by changing their expression levels. Usually, these drought stress-inducible genes have been divided into two groups. One group that directly protects plants against drought stress are involved in water transport (aquaporin) (Alexandersson et al., 2005; 2010), scavenging of free oxygen radicals (superoxide dismutase, catalase, and peroxidase), maintaining cellular membrane integrity (proline, mannitol, glycine, and betaine), and protecting macromolecules (chaperones and late embryogenesis abundant proteins) (Shinozaki et al., 2007; Golldack et al., 2014). The second group is involved in signal perception, signal transduction, and amplification. These include receptor proteins, protein kinases, protein phosphatases, and transcription factors (TFs) (Shinozaki et al., 2007; Golldack et al., 2014). To date, many drought stress-inducible genes, especially transcription factors, have been functionally demonstrated to play crucial roles in plant drought tolerance. These transcription factors include ABA-dependent MYC/MYB and WRKY, ABA-responsive element binding/ABA-binding factor (AREB/ABF), ABA-independent dehydration-responsive element-binding proteins (DREB), C-repeat/drought-responsive element (CRT/DRE), and NAC transcription factors (Shinozaki et al., 2003; Yamaguchi-Shinozaki et al., 2006; Jeong et al., 2010; Ren et al., 2010; Shin et al., 2011; Hu and Xiong, 2014).

Microarray-based analysis has vastly contributed to our understanding of the molecular mechanisms involved in plant drought tolerance. However, there are specific probe design and RNA variant detection constraints (Valdés et al., 2013). As an alternative, RNA sequencing (RNA-seq) technology, which increases specificity and sensitivity, has emerged as a powerful technique for the detection of genes, transcripts, and differential expression profiling. It can be used to monitor gene function at the entire genome level in a species without any available genome information (Wang et al., 2013). RNA-seq technology has been used to dissect the molecular responses of plant drought tolerance in many plants, especially in non-model plants without available genome information, and some new drought stress genes have been identified (Yates et al., 2014; Wu et al., 2014; Li et al., 2015; Bhardwaj et al., 2015; Fu et al., 2016; Li et al., 2016). Although gene microarray and RNA-seq technology have led to major advances in understanding plant responses to drought, knowledge about the molecular mechanisms underlying drought tolerance in medicinal plants is still extremely limited.

*Marsdenia tenacissima* is a well-known anti-cancer medicinal plant used in traditional Chinese medicine. It is widely distributed in tropical to subtropical areas across Asia, particularly in Guizhou and Yunnan Provinces, China (Yu et al., 2011). *Marsdenia tenacissima* can also be used to treat asthma, tracheitis, tonsillitis, pharyngitis, cystitis, and pneumonia (Huang et al., 2013; Li et al., 2013; Deng et al., 2013; Zheng et al., 2014). Drought stress clearly negatively impacts the normal growth and development of*M. tenacissima*, which leads to yield losses and plant quality decline (Meng et al., 2015). However, to date, the molecular mechanism controlling drought tolerance in *M. tenacissima* is unknown, and no drought tolerance gene has been identified. In this study, we performed a comprehensive transcriptome sequencing analysis to explore the drought-tolerance mechanism in *M. tenacissima* and to identify the candidate genes that could potentially be used to improve crop drought resistance.

## Materials and methods

### Plant material, growth conditions, and drought stress treatments

The *M. tenacissima* strain “Yunnan” was used in this study, which was supplied by Yunnan Xintong Plant Pharmaceutical Co., Ltd. (Mengzi,Yunnan,China). The *M. tenacissima* seeds were surface-sterilized in 0.5% (w/v) NaClO for 15 min. Then they were sown in pots filled with peat and vermiculite (v/v = 3:1), and left to germinate in a greenhouse at 25°C. The two-week-old *M. tenacissima* seedlings were individually transferred to a small flowerpot containing 1 kg soil (humus soil:garden soil = 1:1) and grown in an artificial climate incubator under natural drought stress treatment(12 h/12 day/night, light 4000 lx,temperature:23°C/16°C day/night, air relative humidity: 75%/55% day/night).

In the drought treatment, 10–15-cm high plants were split into three groups with ten plants in each group. The first group of plants was supplied water every two days as the control (CK). The second group of plants was not supplied water until the plant drought phenotype (T1) appeared. The last group of plants was not supplied water until the plant drought phenotype appeared (the degree of drought was the same as T1), then sample were taken after watering for 24 hours (T2).The roots, stems, and leaves from three randomly selected plants in each group were collected and stored at −80°C for RNA extraction.

### Total RNA extraction and cDNA synthesis and sequencing

Total RNA was extracted from each sample using Triazol reagent (TaKaRa, Dalian, China) according to the manufacturer’s instructions. The samples were then treated with DNase I to remove any contaminated genomic DNA. The integrity and purity of the RNA was verified by an ultraviolet spectrophotometer (OD260/OD280 ratios of 1.89 to 2.08) and 1.2% agarose gel electrophoresis. The RNA from the roots, stems, and leaves of each group of plants was pooled. The cDNA libraries were then constructed according to Huang et al. (2014). The cDNA libraries were sequenced on a HiSeq2000 (Illumina, San Diego, CA, USA) according to the manufacturer’s standard protocols to generate 100-bp paired-end reads.

### Acquisition of clean reads and mapping

Raw reads from the cDNA library were filtered to remove low-quality reads and adaptors using the program FASTX-Tool kit (http://hannonlab.cshl.edu/fastx_toolkit/) to produce the clean reads. The clean reads were mapped to the reference transcriptome dataset (NCBISRA140234) using SOAP aligner/soap2 software (Li et al., 2009). The total mapped reads were kept for further analysis.

### Identification of differentially expressed genes (DEGs)

The DEGs between treatments were identified based on the Reads Per Kilobase per Million (RPKM) value calibrated by DEGseq (Gong et al., 2015; Wang et al., 2010). Genes with a “q value < 0.005” and a “fold change |log2| > 1” were deemed to be significantly differentially expressed between the two samples.

### Functional annotation and classification

The DEGs were annotated using the following databases: the NR protein database (NCBI), Swiss Prot, Gene Ontology (GO), the Kyoto Encyclopedia of Genes and Genomes (KEGG) database, and the Clusters of Orthologous Groups database (COG) according to the methods of described by Zhou et al. (2014). Pathways and GO function enrichment analyses were performed as previously described (Sun et al., 2013). The transcription factor (TF) responses to drought stress were identified according to the method described by Zhao et al. (2016).

### Quantitative real-time PCR (qRT-PCR) verification of DEGs

To evaluate the accuracy and validity of the transcriptome sequencing data, 24 genes with differential expressions were selected to carry out the qRT-PCR analysis. Primers were designed using the BioXM 2.6 software, and the primer sequences are listed in Table S1. The GAPDH (glyceraldehyde-3-phosphate dehydrogenase) gene was used as a reference gene. The qRT-PCR analysis of each gene was performed with three biological replicates according to the SYBR Premix ExTaq™ protocol (TaKaRa) on a Light Cycler 480 Real-Time PCR machine (Roche Diagnostics Ltd., Switzerland). The relative expression level of each gene was calculated using the 2^-(ΔΔCt)^ method. The expression value of each gene from qRT-PCR and RNA-seq was log2 transformed so that the qRT-PCR data could be compared with the RNA-seq results.

## Results

### Transcriptome sequencing, data statistics and evaluation, and reads mapping

To understand the drought-response molecular mechanism in *M. tenacissima* and identify potential candidate genes involved in drought tolerance, deep RNA sequencing of*M. tenacissima* seedlings subjected to drought and subsequent rewatering was performed using the Illumina sequencing platform. A total of 43,983,844, 48,059,552, and 43,744,500 raw reads were obtained from the CK, T1, and T2 cDNA libraries, respectively (Table 1). After removing the low-quality reads and adaptors, 43,129,228, 47,116,844, and 42,815,454 clean reads were produced, which accounted for 98.06%, 98.04%, and 97.88% of the raw reads, respectively (Table 1). Furthermore, 32,879,580 (76.24 %), 36,085,718 (76.59%), and 34,482,932 (80.54%) clean reads were mapped to the reference transcriptome (NCBI SRA140234) and 44,112, 39,307, and 39,608 genes were generated by SOAP aligner/soap2 software, respectively (Table 1). The total mapped reads were used to estimate the gene expression levels.

**Table 1.** Original data statistics.

| Analysis reads | CK | T1 | T2 |
| --- | --- | --- | --- |
| Raw reads | 43,983,844 | 48,059,552 | 43,744,500 |
| Clean reads | 43,129,228 | 47,116,844 | 42,815,454 |
| Clean bases (Gb) | 5.02 | 5.48 | 4.98 |
| Q20 | 98.06 | 98.04 | 97.88 |
| Average length | 859 | 985 | 975 |
| Total mapped reads (%) | 32,879,580<br>(76.24) | 36,085,718<br>(76.59) | 34,482,932<br>(80.54) |
| Unique mapped reads (%) | 32,825,505<br>(76.11) | 36,000,522<br>(76.41) | 34,428,760<br>(80.41) |
| Gene number | 44,112 | 39,306 | 39,607 |

### Identification of DEGs responding to drought stress

The genes from each treatment group were subjected to a pairwise comparison to identify the DEGs after using a blast algorithm with the preset cutoffs. As a result, a total of 21,254 DEGs were identified. A comparison between CK and T1 showed that 1855 genes were up-regulated, and 6817 genes were down-regulated. Between CK and T2, 1612 genes were up-regulated and 4431 genes down-regulated. A further 3982 genes were found to be up-regulated, and 2555 genes were down-regulated between T1 and T2, and 78 up-regulated genes and 144 down-regulated genes were identified in all three comparison groups (Fig. 1). Notably, 1174 of the 1855 induced genes and 2344 of the 6817 repressed genes under drought stress were down-regulated, but were up-regulated after re-watering (Fig. 2, Tables S1–S2). These genes probably play important roles in tolerance to drought stress.

**Figure 1.**
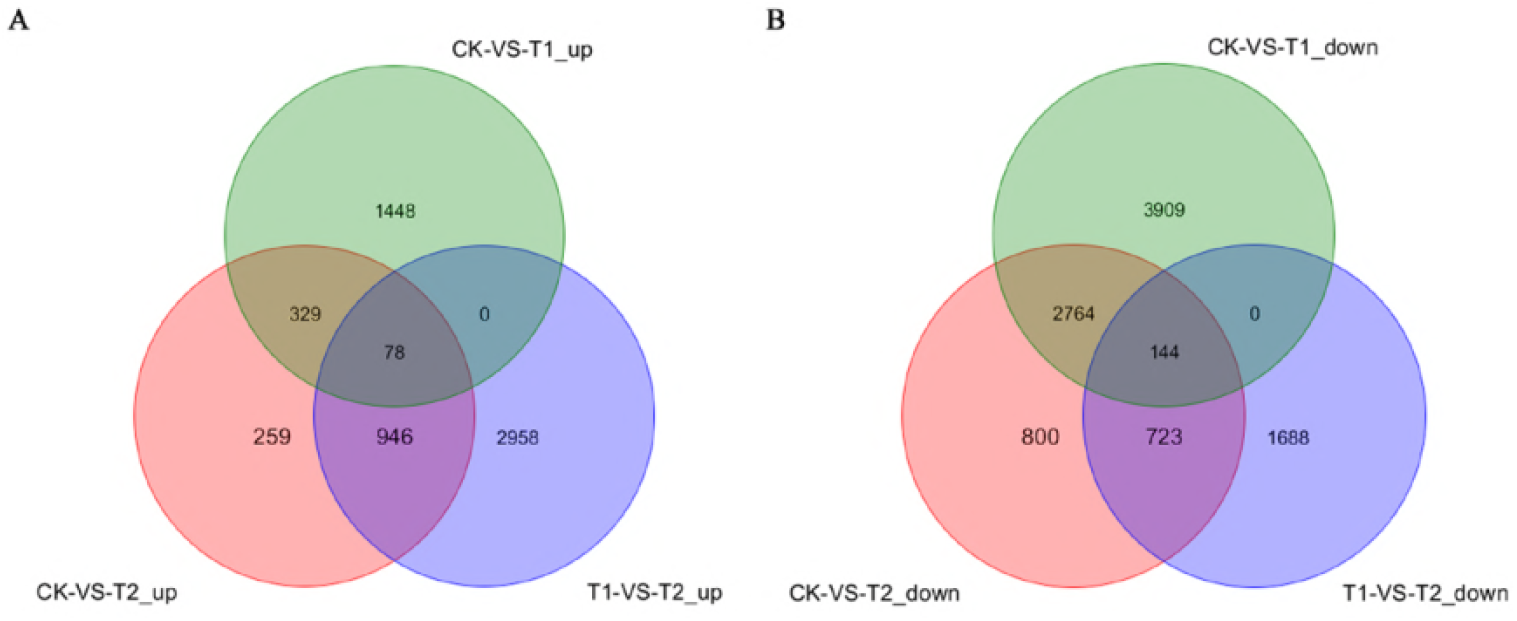
Venn diagram analysis of differentially expressed genes. The numbers of differentially expressed genes are shown in the diagram; CK – control (no treatment); T1 – drought stress; T2 – re-watering treatment.

**Figure 2.**
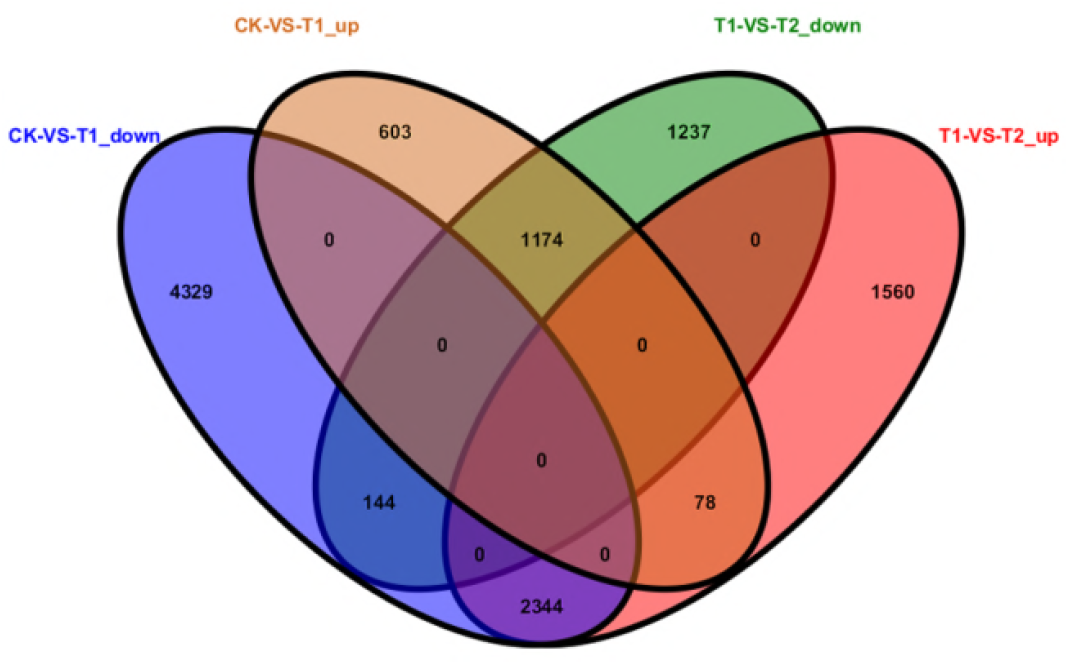
Venn diagram analyses of differentially expressed genes between CK vs. T1, and T1 vs. T2.

### GO annotation of DEGs from the three comparison groups

GO classification was performed to investigate the functions of the DEGs in the three comparison groups. After comparing CK with T1, 2628 DEGs (704 up-regulated genes and 1924 down-regulated genes) were assigned to 67 main functional groups in the “biological processes”, “cellular components”, and “molecular functions” categories. When CK was compared to T2, 1699 DEGs (559 up-regulated genes and 1140 down-regulated genes) could be functionally assigned to the relevant terms. The T1 versus T2 comparison functionally assigned 2161 DEGs (1296 up-regulated genes and 865 down-regulated genes) to the relevant terms. The top three significantly enriched GO functional annotation categories were “metabolic process”, “cell”, and “catalytic activity”(Fig. 3).

**Figure 3.**
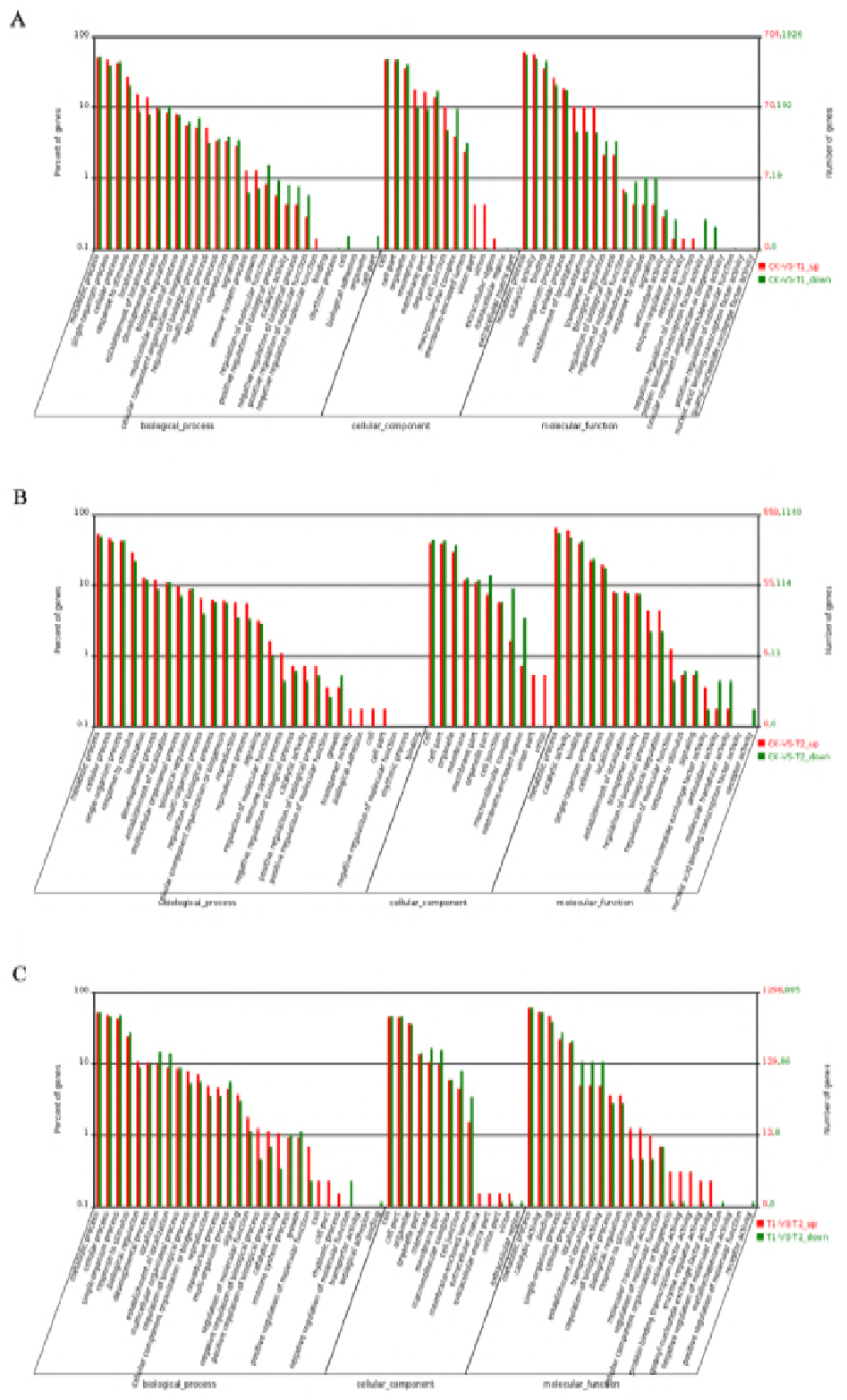
Gene Ontology (GO) categorization of the differentially expressed genes in the three comparison groups. A, CK vs. T1; B, CK vs. T2; C, T1 vs. T2

### KEGG pathway analysis of DEGs

To determine whether these DEGs engaged in specific pathways, we performed a detailed KEGG pathway classification by searching against the KEGG pathway database. A total of 1910 of the DEGs from CK vs. T1 could be annotated into 120 pathways. The top five pathways were “metabolic pathways” (553), “biosynthesis of secondary metabolites” (229), “ribosome” (143), “starch and sucrose metabolism” (71), and “oxidative phosphorylation” (60) (Table S3). For CK vs. T2, a total of 1210 of the DEGs could be classified into 114 pathways. The top five pathways were “metabolic pathways” (352), “biosynthesis of secondary metabolites” (205), “ribosome” (92), “plant hormone signal transduction” (47), and “oxidative phosphorylation” (47) (Table S4). For T1 vs. T2, a total of 1426 of the DEGs could be annotated into 115 pathways. The top five pathways were “metabolic pathways” (457), “biosynthesis of secondary metabolites” (255), “ribosome” (65), “starch and sucrose metabolism” (63), and “plant hormone signal transduction” (58) (Table S5).

### Transcription factor responses to drought stress and water stimulus

Transcription factors are known to play vital roles in plant abiotic stress tolerance because they can regulate the expression of numerous downstream genes. A total number of 1039, 1016, and 980 TFs were identified in CK, T1, and T2, respectively (Table 2). The number of TFs identified in the T1 and T2 library was slightly less than in CK. In addition, 363, 267, and 299 TFs were identified as DEGs in CK vs. T1, CK vs. T2, and T1 vs. T2, respectively. Further analysis revealed that the 363 DEGs from CK vs. T1 could be grouped into 42 families, and the top five families were C2H2 (35), bHLH (33), ERF (26), NAC (24), and MYB (21) (Fig. 4A, Table S6). Similarly, the 267 DEGs from CK vs T2 were grouped into 37 families, and most of the DEGs (103) belonged to the C2H2, ERF, bHLH, and bZIP families (Fig. 4B, Table S7). Furthermore, in the comparison between T1 and T2, 299 DEGs were involved in a total of 43 TF families. Among these TF families, ERF (37), bHLH (31), MYB (25), C2H2 (19), and MYB-related (18) were the top five families with the most genes (Fig. 4C, Table S8).

**Figure 4.**
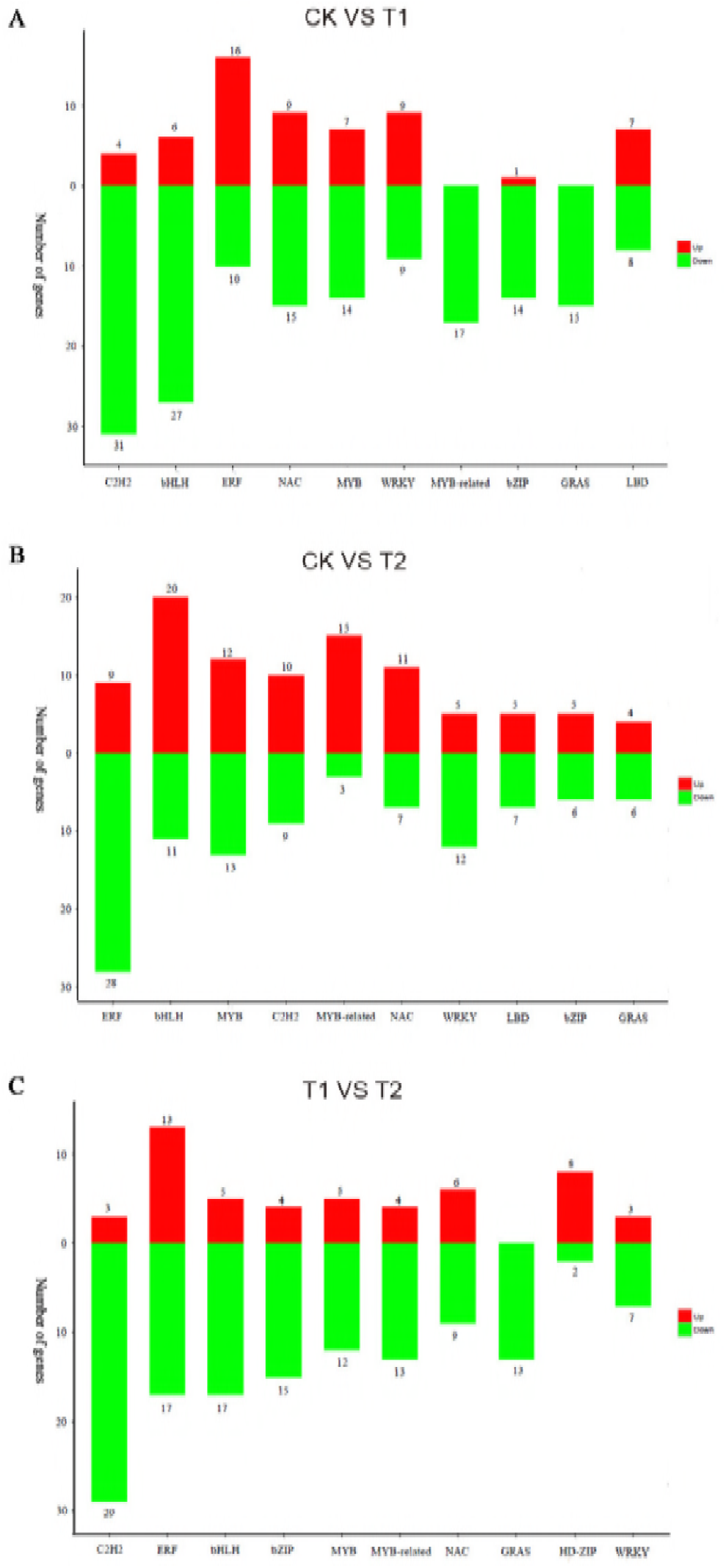
The top 10 families of differentially expressed transcription factors in the CK vs. T1 group (A), the CK vs. T2 group (B), and the T1 vs. T2 group (C).

**Table 2.** Transcription factors (TFs) in the sequencing libraries.

| TFs | CK vs T1 | CK vs T2 | T1vs T2 |
| --- | --- | --- | --- |
| Total differentially expressed TFs | 363 | 267 | 299 |
| Up-regulated differentially expressed TFs | 81 | 77 | 174 |
| Down-regulated differentially expressed TFs | 282 | 190 | 125 |

A further analysis of the quantity relationship between up- and down-regulated TFs in the three comparison groups found that the number of up-regulated genes was significantly lower than the number of down-regulated genes in both CK vs. T1 and CK vs. T2, but was the reverse in T1 vs. T2 (Table 2). In the CK vs. T1 comparison, the number of down-regulated TFs was 3-fold more than the up-regulated ones. The largest number of down-regulated TFs was found in the C2H2 family, while the ERF family contained the largest number of up-regulated TFs (Fig. 4A). Similarly, in CK vs. T2, the number of down-regulated TFs was 2-fold higher than the up-regulated ones. The ERF and bHLH families contained the largest number of down-regulated and up-regulated TFs, respectively (Fig. 4B). In T1 vs. T2, the number of down-regulated TFs was lower than the number of up-regulated TFs. The C2H2 family contained the largest number of down-regulated TFs, whereas the ERF family had the largest number of up-regulated TFs (Fig. 4C).

### Identification of candidate genes for drought stress resistance

To identify the candidate genes for drought stress resistance in *M. tenacissima*, 1174 genes that were induced by drought stress and repressed by re-watering were screened. A blast analysis showed that 855 of the 1174 genes had a functional description (Table S9). Further analysis found that 64 genes were homologous to the known functional genes that directly protects plants against drought stress, which include aquaporin, late embryogenesis abundant protein, chaperone, dehydration responsive protein, pleiotropic drug resistance protein, alcohol dehydrogenase, peroxidase, proline metabolism genes, trehalose synthesis-related genes, flavonoid synthesis-related genes, mannitol transporter, sugar transporter, peptide transporter, MATE efflux protein, and ABC transporter genes (Table 3). In addition, histone, histone deacetylase, and methyltransferase were also found, which suggested that epigenetic regulation was involved in the *M. tenacissima* drought stress resistance mechanism (Table 3).

**Table 3.** The putative functional genes that were induced by drought stress and repressed by re-watering.

| Gene ID | Fold change (T1/CK) | Fold change (T2/T1) | Blast swissprot |
| --- | --- | --- | --- |
| Unigene0008635 | 3.1 | -2.2 | glutamate dehydrogenase A-like [Solanum lycopersicum] (Arginine and proline metabolism) |
| Unigene0012721 | 10.9 | -4.8 | trehalose-6-phosphate synthase [Camellia sinensis] |
| Unigene0045759 | 2.3 | -4.2 | probable trehalose-phosphate phosphatase J-like [Solanum lycopersicum] |
| Unigene0022077 | 8.5 | -3.7 | alpha-trehalose-phosphate synthase [UDP-forming] 9-like [Solanum lycopersicum] |
| Unigene0027014 | 2.1 | -16.6 | Late embryogenesis abundant hydroxyproline-rich glycofamily protein [Theobroma cacao] |
| Unigene0009110 | 11.9 | -8.1 | Late embryogenesis abundant hydroxyproline-rich glycofamily protein [Theobroma cacao] |
| Unigene0013711 | 3.8 | -7.9 | late embryogenesis abundant protein group 9 protein, partial [Genlisea aurea] |
| Unigene0023521 | 2.5 | -3.3 | aquaporin 1 [Nicotiana tabacum] |
| Unigene0021777 | 2.2 | -3.0 | aquaporin protein AQU20 [Camellia sinensis] |
| Unigene0009422 | 14.5 | -15.4 | dehydration responsive protein [Corchorus olitorius] |
| Unigene0019806 | 2.3 | -5.7 | dehydration responsive protein [Corchorus olitorius] |
| Unigene0020350 | 2.0 | -9.6 | probable mitochondrial chaperone BCS1-B-like [Fragaria vesca subsp. vesca] |
| Unigene0016740 | 4.7 | -7.0 | chaperone protein dnaJ 11, chloroplastic-like [Vitis vinifera] |
| Unigene0012659 | 3.1 | -2.2 | Heat shock protein 70 (Hsp 70) family protein [Theobroma cacao] |
| **Unigene0016751** | 2.5 | -2.4 | Heat shock protein DnaJ with tetratricopeptide repeat isoform 1 [Theobroma cacao] |
| **Unigene0008874** | 3.1 | -4.1 | copper chaperone [Populus alba x Populus glandulosa] |
| **Unigene0022185** | 2.8 | -2.2 | pleiotropic drug resistance protein 1-like [Vitis vinifera] |
| **Unigene0024248** | 13.8 | -11.5 | pleiotropic drug resistance protein 2-like [Vitis vinifera] |
| **Unigene0026840** | 3.2 | -2.5 | pleiotropic drug resistance protein 2-like [Solanum lycopersicum] |
| **Unigene0010544** | 5.2 | -5.1 | alcohol dehydrogenase-like protein [Ocimum basilicum] |
| **Unigene0014656** | 2.2 | -2.7 | alcohol dehydrogenase [Solanum tuberosum] |
| **Unigene0032109** | 25.4 | -22.2 | cinnamyl alcohol dehydrogenase [Neolamarckia cadamba] |
| **Unigene0021885** | 158.8 | -49.6 | sinapyl alcohol dehydrogenase-like 3 [Nicotiana tabacum] |
| **Unigene0026625** | 2.1 | -5.5 | peroxidase 4 [Vitis vinifera] |
| **Unigene0007407** | 4.8 | -12.5 | peroxidase 10-like [Solanum lycopersicum] |
| **Unigene0037539** | 6.4 | -4.5 | peroxidase [Populus alba x Populus glandulosa] |
| **Unigene0020714** | 3.0 | -4.4 | peroxidase 4 [Litchi chinensis] |
| **Unigene0008948** | 5.2 | -4.6 | peroxidase 25 [Vitis vinifera] |
| **Unigene0045735** | 3.8 | -6.1 | peroxidase 73-like [Solanum lycopersicum] |
| **Unigene0023720** | 3.2 | -4.3 | phenylalanine ammonia lyase [Catharanthus roseus] |
| **Unigene0023719** | 3.8 | -3.4 | phenylalanine ammonia lyase [Catharanthus roseus] |
| **Unigene0022294** | 3.0 | -2.7 | hydroxycinnamoyl-CoA quinate hydroxycinnamoyl transferase [Coffea canephora] |
| **Unigene0021581** | 12.1 | -2.6 | flavonoid 3'-monooxygenase-like [Solanum lycopersicum] |
| **Unigene0017542** | 2.4 | -2.1 | hydroxycinnamoyl-CoA shikimate/quinate hydroxycinnamoyl transferase [Coffea canephora] |
| **Unigene0018052** | 5.0 | -6.3 | mannitol transporter [Artemisia annua] |
| **Unigene0023659** | 2.1 | -2.9 | mannitol transporter [Artemisia annua] |
| **Unigene0019188** | 2.7 | -2.6 | sugar transport protein [Coffea canephora] |
| **Unigene0023657** | 3.7 | -2.5 | sugar transport protein [Coffea canephora] |
| **Unigene0021497** | 8.1 | -55 | sugar transporter ERD6-like 7-like [Vitis vinifera] |
| **Unigene0014067** | 3.6 | -6.5 | sugar transport protein 14-like [Solanum lycopersicum] |
| **Unigene0036310** | 16.1 | -17.0 | bidirectional sugar transporter SWEET16-like [Solanum lycopersicum] |
| **Unigene0041988** | 21.5 | -45.1 | sugar transporter ERD6-like 16-like [Solanum lycopersicum] |
| **Unigene0023558** | 2.1 | -2.6 | bidirectional sugar transporter SWEET2a-like [Solanum lycopersicum] |
| **Unigene0041404** | 4.2 | -7.7 | bidirectional sugar transporter NEC1-like [Solanum lycopersicum] |
| **Unigene0015567** | 5.1 | -5.3 | peptide transporter PTR3-A-like [Fragaria vesca subsp. vesca] |
| **Unigene0021742** | 8.9 | -7.0 | Peptide transporter 1 isoform 1 [Theobroma cacao] |
| **Unigene0015568** | 2.4 | -2.4 | peptide transporter PTR3-A-like [Solanum lycopersicum] |
| **Unigene0021745** | 8.1 | -8.2 | Peptide transporter 1 isoform 1 [Theobroma cacao] |
| **Unigene0015566** | 3.1 | -3.7 | peptide transporter PTR3-A-like [Cucumis sativus] |
| **Unigene0023591** | 5.1 | -4.4 | MATE efflux family protein 5 [Vitis vinifera] |
| **Unigene0014060** | 3.1 | -2.1 | MATE efflux family protein [Theobroma cacao] |
| **Unigene0007370** | 5.6 | -7.9 | MATE efflux family protein [Theobroma cacao] |
| **Unigene0023504** | 6.0 | -3.6 | MATE efflux family protein [Theobroma cacao] |
| **Unigene0031906** | 4.9 | -3.5 | MATE efflux family protein 8-like [Fragaria vesca subsp. vesca] |
| **Unigene0018290** | 4.4 | -2.5 | ABC transporter C family member 4-like [Solanum lycopersicum] |
| **Unigene0023528** | 2.4 | -4.8 | ABC transporter B family member 21-like [Solanum lycopersicum] |
| **Unigene0035979** | 11.7 | -3.2 | mutant histone deacetylase 6 [Arabidopsis thaliana] |
| **Unigene0006009** | 2.1 | -3.5 | histone-lysine N-methyltransferase ASHR2-like isoform 1 [Solanum lycopersicum] |
| **Unigene0012248** | 2.1 | -3.3 | Histone H3 K4-specific methyltransferase SET7/9 family protein [Theobroma cacao] |
| **Unigene0009759** | 2.0 | -3.9 | Methyltransferases [Theobroma cacao] |
| **Unigene0013388** | 2.1 | -3.3 | histone H1 [Solanum lycopersicum] |
| **Unigene0012434** | 3.5 | -7.9 | Histone superfamily protein [Theobroma cacao] |
| **Unigene0007578** | 2.1 | -3.7 | Histone H2B [Medicago truncatula] |
| **Unigene0043003** | 3.0 | -4.6 | PREDICTED: histone H2AX [Vitis vinifera] |

To further identify the crucial regulatory genes, we investigated the protein kinases and the transcription factors in the 855 genes with functional descriptions. A total of 44 protein kinases were identified, which could be classified into 13 class-types. The top four classes were receptor-like protein kinase (11), L-type lectin-domain containing receptor kinase (6), LRR receptor-like serine/threonine-protein kinase (5), and leucine-rich repeat receptor-like protein kinase (5) (Table 4). A total of 38 transcription factors were identified, which could be categorized into eight TF families. Among these TF families, ERF (10), WRKY (8), and NAC (5) were the top three families with the most genes (Table 4).

**Table 4.**
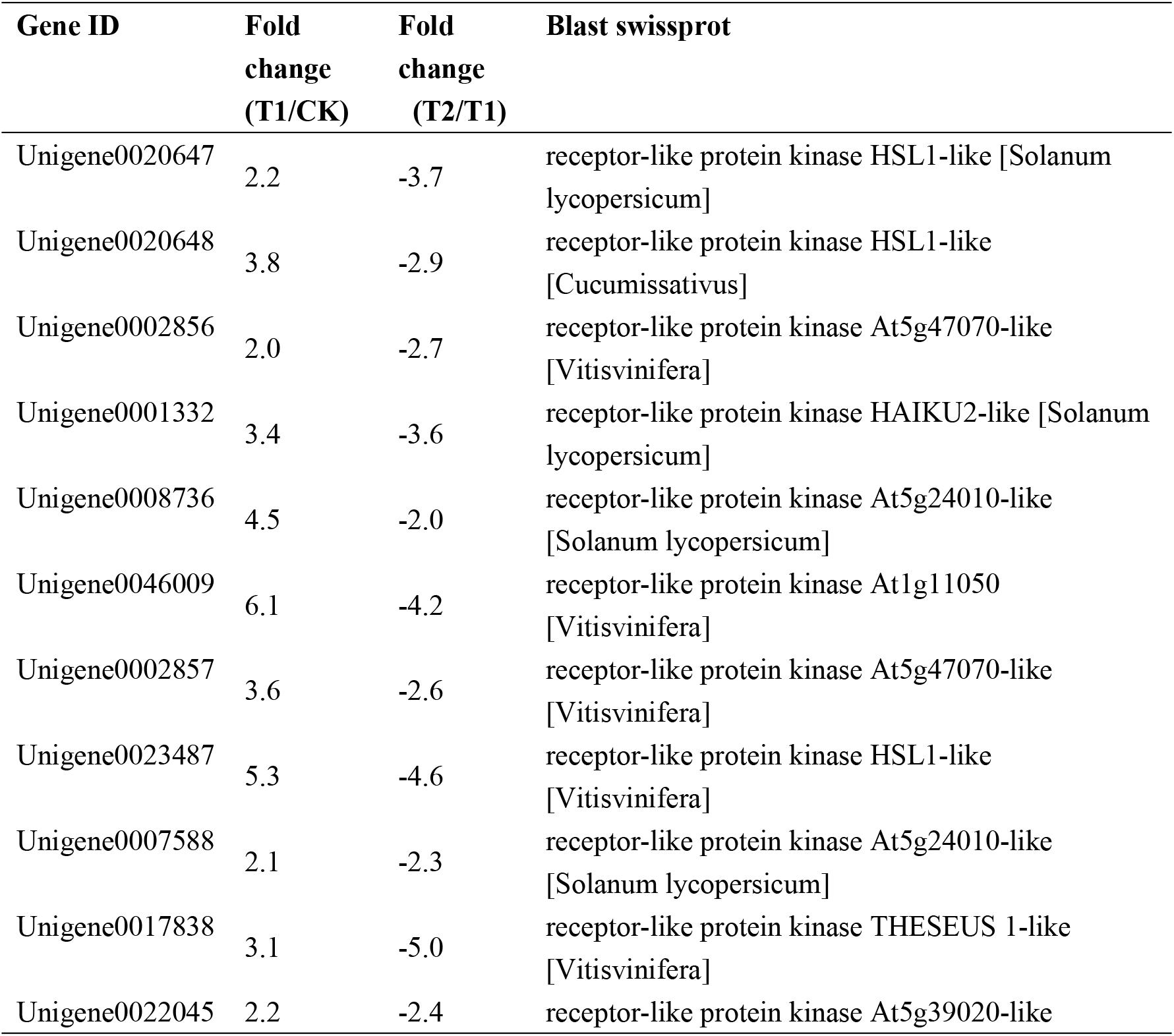
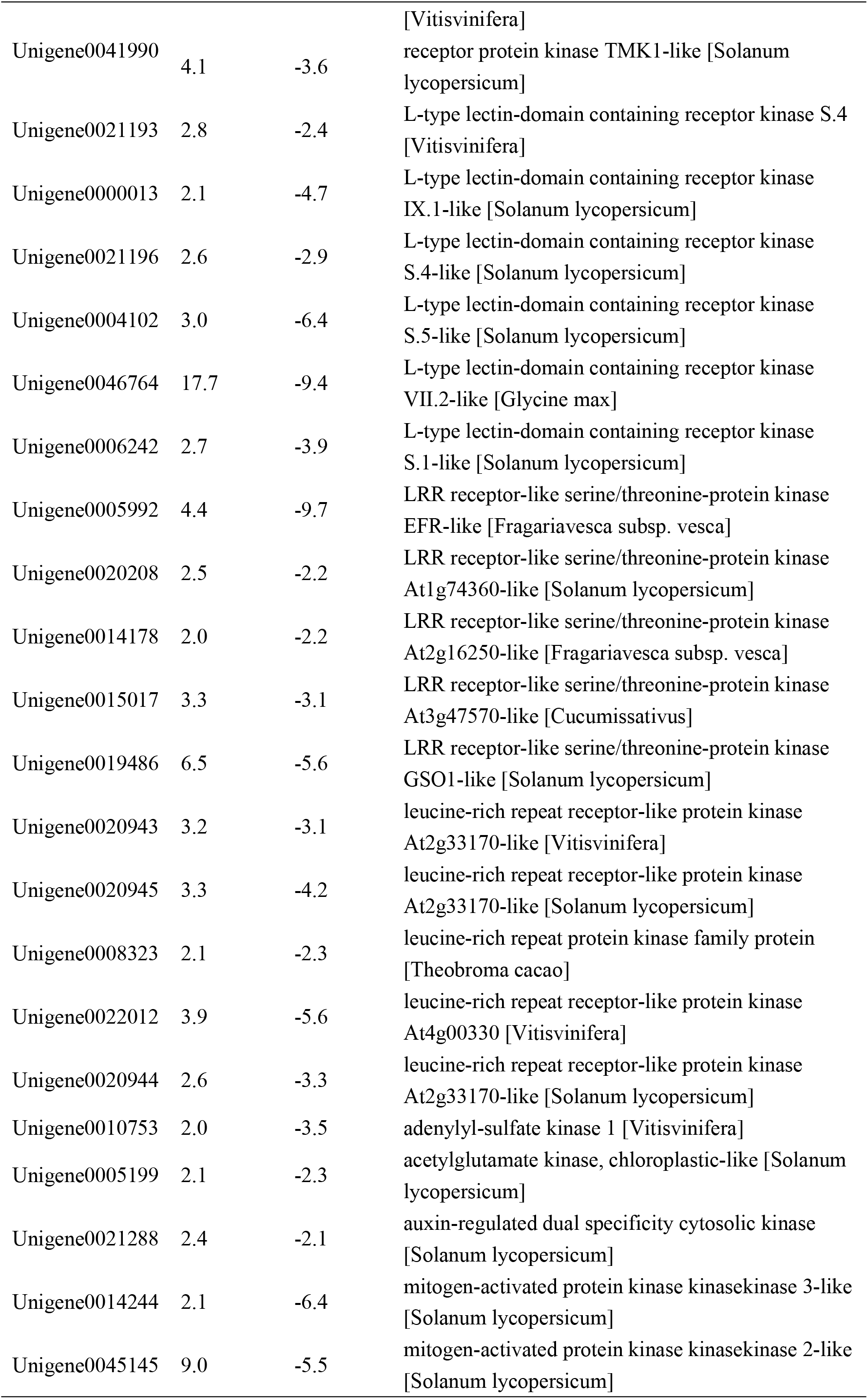
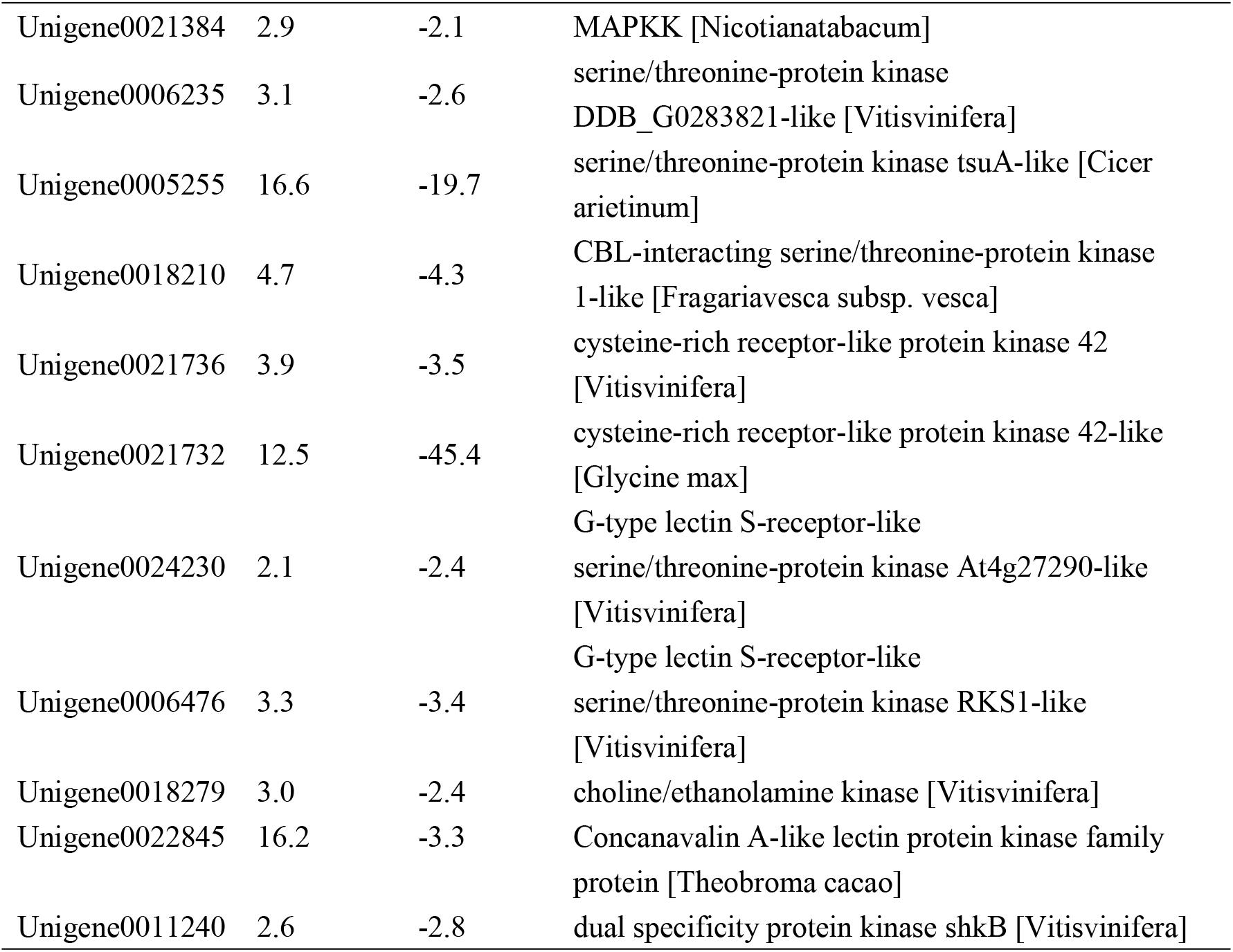
The putative kinase encoding genes that were induced by drought stress and repressed by re-watering.

### Validation of the RNA-seq data by qRT-PCR

To verify the validity of the RNA-seq data, we analyzed 24 genes using qRT-PCR (Table S10). The correlation coefficients of the gene expression trends after qRT-PCR, and the sequencing data from CK vs. T1and CK vs. T2 were 0.6315 and 0.5735, respectively (Fig. 5), which confirmed the validity of the RNA-seq data.

**Figure 5.**
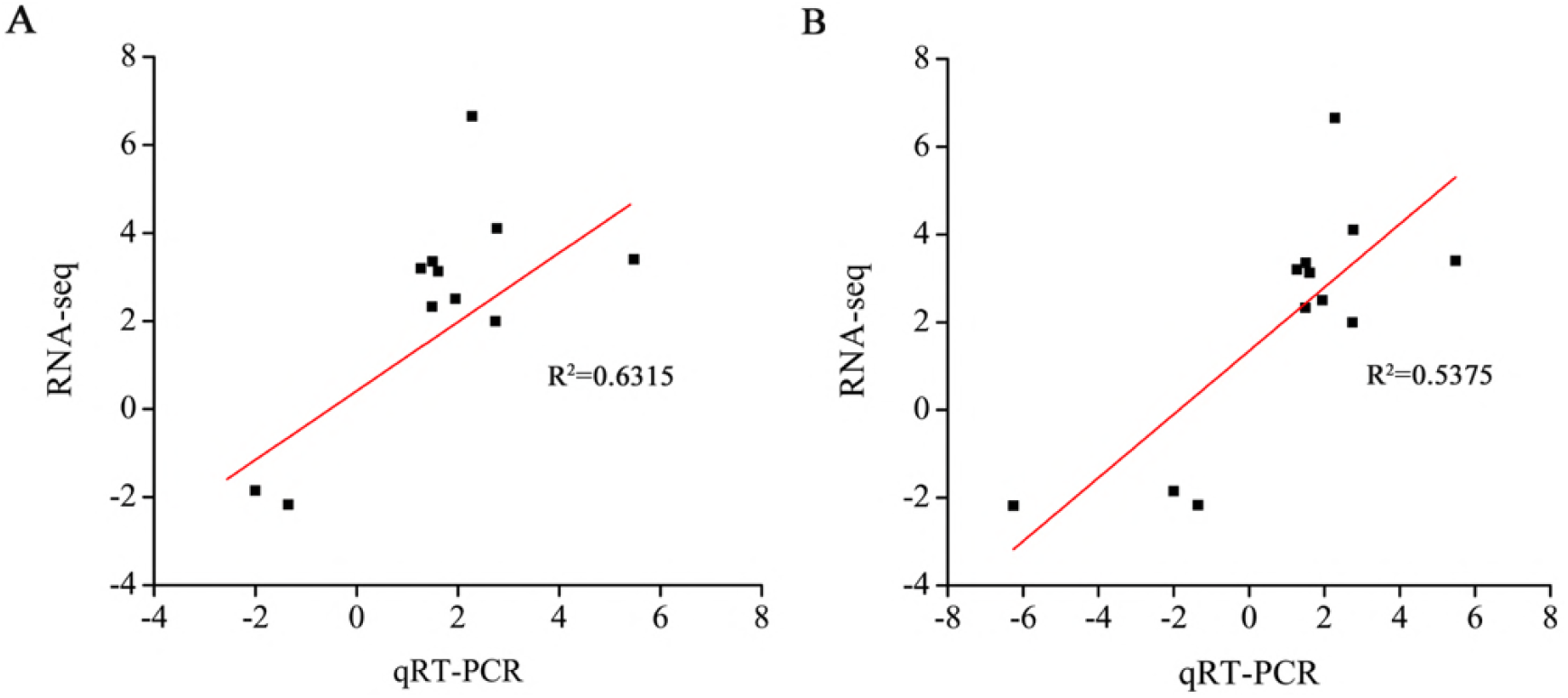
Verification of the differentially expressed genes in the CK vs .T1 group (A) and the CK vs. T2 group (B) by qRT-PCR.

## Discussion

In this study, two aquaporin genes were up-regulated by drought stress and down-regulated after re-watering. This indicated that these aquaporin(s) may play a similar regulatory role in *M. tenacissima* under drought stress. The results have been confirmed in chickpea, foxtail millet, maize, and potato (Hayano-Kanashiro et al., 2009; Jain and Chattopadhyay, 2010; Lata et al., 2010; Gong et al., 2015). Heat shock responsive genes are functional genes that facilitate protein refolding and stabilize polypeptides and membranes. They have also been reported to respond to drought stress in barley, rice, and potato (Rabbani et al., 2003; Talamè et al., 2007; Gong et al., 2015). In this study, we found that the expression of two heat shock responsive genes were induced by drought stress and repressed by re-watering treatment, which suggested that they directly participated in regulating drought-stress responses in *M. tenacissima*. Generally, drought stress induces the accumulation of LEA proteins and this accumulation enhances the survival rate of plants under drought conditions (Borovskii et al., 2002; Jiang and Huang, 2002; Porcel et al., 2005; Guo et al., 2009; Liu and Jiang, 2010; Zhao et al., 2016). Some studies have suggested that the role of LEA proteins was to facilitate the correct folding of both structural and functional proteins and prevent lipid peroxidation (Liu and Jiang, 2010). In this study, the expression of three LEA encoding genes was significantly up-regulated by drought stress and down-regulated by re-watering treatment, which suggested that these genes were involved in drought stress resistance in *M. tenacissima*.

Trehalose has a protective role against various abiotic stresses, including drought stress in bacteria, fungi, and some plants. It helps maintain cellular membrane integrity and prevent protein degradation (Seki et al., 2007; Delorge et al., 2014). In plants, trehalose-6-phosphate synthase (TPS) and trehalose-6-phosphate phosphatase catalyze the biosynthesis of trehalose, and their expressions are induced by drought stress (Avonce et al., 2004; Paul, 2007; Li et al., 2011; Zhao et al., 2016).

Furthermore, over expression *AtTPS1* or *OsTTPS1* improved the stress resistance of transgenic plants (Avonce et al., 2004; Paul, 2007; Li et al., 2011). In this study, one TPP gene and two TPS encoding genes were up-regulated by drought stress and down-regulated after re-watering. Furthermore, one proline synthesis-related gene had a similar expression pattern to the TPP and TPS genes. These results indicate that synthesizing compatible solutes is a conserved drought resistance mechanism in different plants.

It well-known that drought stress produces reactive oxygen species (ROS) and excessive ROS can cause the irreversible oxidization of lipids and proteins, which leads to membrane injury (Li et al., 2012). To overcome ROS injury, plants utilize ROS-scavenging enzymes, such as peroxidase (POD), superoxide dismutase (SOD), and catalase (CAT), to scavenge the excessive ROS (Koussevitzky et al., 2008). In this study, we found that six POD-encoding genes were significantly up-regulated by drought stress and down-regulated by re-watering treatment. This result suggested that ROS-scavenging via POD is an important mechanism in the overall resistance of *M. tenacissima* to drought stress.

Some studies have shown that alcohol dehydrogenase, MATE efflux family protein, and ABC transporters are involved in protecting plants against drought stress (Klein et al., 2004; Xiao et al., 2009; Kuromori et al., 2011; Zhang et al., 2014; Zhang et al., 2017). In this study, three alcohol dehydrogenase-, five MATE-, and two ABC-encoding genes were identified, and their expressions were strongly induced by drought stress, but significantly repressed by re-watering treatment, which suggested that these genes may also play important roles in drought stress resistance.

Many TFs, such as ABF, bHLH, ERF, MYB, NAC, and WRKY, act as key regulators and play crucial roles in plant resistance to drought stress because they regulate many downstream functional genes (Shinozaki et al., 2003; Yamaguchi-Shinozaki et al., 2006; Jeong et al., 2010; Ren et al., 2010; Shin et al., 2011; Hu and Xiong, 2014). This study identified 38 TFs that were significantly up-regulated by drought stress and down-regulated after re-watering. The 38 TFs could be classified into the AP2/ERF, bHLH, BES1, ERF, MYB, MYB-related, NAC, WRKY, and Trihelix families. Among these nine families, the ERF (10), WRKY (8), NAC (5), and AP2/ERF (4) families accounted for 70% of the genes. This indicated that these TFs may have important functions in regulating resistance to drought stress and can be used as candidate genes to further investigate drought stress in *M. tenacissima*.

## Conclusion

In this study, we performed a comparative analysis of the transcriptome changes in *M. tenacissima* undergoing drought stress and re-watering treatment. A total of 8672, 6043, and 6537 DEGs, including 363, 267, and 299 TFs, were identified in the CK vs. T1, CK vs. T2, and T1 vs. T2 comparisons, respectively. The DEGs from these three comparative groups were classified into 67, 58, and 66 GO categories and were involved in 120, 114, and 115 KEGG pathways, respectively. Interestingly, 1174 up-regulated and 2344 down-regulated genes under drought stress had the opposite expression pattern after re-watering. Analysis of the 1174 up-regulated genes induced by drought stress and repressed by re-watering showed that many genes were homologous to known functional genes that directly protect plants against drought stress. Furthermore, 44 protein kinases and 38 TFs with opposite expression patterns under drought stress and re-watering were identified as crucial candidate regulators of drought stress resistance in *M. tenacissima*.

In summary, our study is the first to characterize the *M. tenacissima* transcriptome in response to drought stress, and has identified the key candidate drought stress resistant genes in *M. tenacissima*. Our results will help unravel the mechanism controlling *M. tenacissima* drought stress resistance.

**Table 5.** The putative TFs genes that were induced by drought stress and repressed by re-watering.

| Gene ID | Fold<br>change<br>(T1/CK) | Fold<br>change<br>(T2/T1) | Blast swissprot |
| --- | --- | --- | --- |
| Unigene0042229 | 27.8 | -13.9 | Ethylene-responsive transcription factor<br>ERF098-like [Solanum lycopersicum] |
| Unigene0004507 | 545.9 | -8.9 | Ethylene-responsive transcription factor ABR1-like<br>[Solanum lycopersicum] |
| Unigene0023064 | 3.5 | -22.2 | Ethylene-responsive transcription factor<br>ERF014-like [Cicer arietinum] |
| Unigene0020372 | 2.7 | -2.6 | Ethylene-responsive transcription factor<br>ERF114-like [Vitisvinifera] |
| Unigene0025466 | 11.8 | -142.9 | Ethylene-responsive transcription factor<br>ERF098-like [Solanum lycopersicum] |
| Unigene0003028 | 3.5 | -12.1 | Ethylene response factor 10 [Actinidiadeliciosa] |
| Unigene0002715 | 2.6 | -6.6 | Ethylene-responsive transcription factor 1A-like<br>[Glycine max] |
| Unigene0023911 | 5.8 | -5.9 | Ethylene response factor 14 [Actinidiadeliciosa] |
| Unigene0003821 | 16.8 | -6.7 | Ethylene response factor 10 [Actinidiadeliciosa] |
| Unigene0017954 | 2.2 | -4.8 | Ethylene-responsive transcription factor RAP2-3 [Vitisvinifera] |
| Unigene0004183 | 2.5 | -2.3 | WRKY transcription factor 53 [Jatropha curcas] |
| Unigene0011852 | 3.4 | -2.9 | WRKY transcription factor 70-like [Vitisvinifera] |
| Unigene0003104 | 3.4 | -2.2 | WRKY transcription factor 48 [Vitisvinifera] |
| Unigene0003461 | 5.5 | -3.3 | WRKY22 [Catharanthusroseus] |
| Unigene0003103 | 2.2 | -2.0 | WRKY transcription factor 31 [(Populustomentosa x P. bolleana) x P. tomentosa] |
| Unigene0013608 | 3.0 | -3.8 | WRKY DNA-binding protein 27 [Arabidopsis thaliana] |
| Unigene0014567 | 2.1 | -2.0 | WRKY transcription factor 22-like [Solanum lycopersicum] |
| Unigene0017815 | 10.7 | -5.5 | WRKY transcription factor 75-like [Setariaitalica] |
| Unigene0012446 | 14.6 | -3.6 | NAC domain containing protein 80 [Theobroma cacao] |
| Unigene0021188 | 2.4 | -3.1 | NAC transcription factor [Camellia sinensis] |
| Unigene0002781 | 3.3 | -3.2 | NAC domain transcriptional regulator superfamily protein [Theobroma cacao] |
| Unigene0006055 | 5.6 | -4.4 | NAC transcription factor 29-like [Solanum lycopersicum] |
| Unigene0012447 | 11.0 | -5.7 | NAC domain-containing protein 100-like [Vitisvinifera] |
| Unigene0036849 | 2.3 | -2.2 | Myb-related protein 2 [Nicotianatabacum] |
| Unigene0030322 | 15.1 | -21.9 | Myb-related protein Myb4-like [Solanum lycopersicum] |
| Unigene0016812 | 4.2 | -3.4 | MYB1 [Gossypiumhirsutum] |
| Unigene0034565 | 4.3 | -4.8 | transcriptional activator Myb-like [Cucumissativus] |
| Unigene0022511 | 12.0 | -4.1 | Myb domain protein 112 isoform 1 [Theobroma cacao] |
| Unigene0000917 | 8.1 | -5.6 | transcription factor MYB75-like [Vitisvinifera] |
| Unigene0016518 | 3.6 | -3.1 | AP2/ERF domain-containing transcription factor [Populustrichocarpa] |
| Unigene0001595 | 6.5 | -8.7 | AP2/ERF domain-containing transcription factor [Populustrichocarpa] |
| Unigene0004302 | 9.8 | -3.5 | AP2/ERF domain-containing transcription factor [Populustrichocarpa] |
| Unigene0046101 | 32.2 | -10.5 | AP2/ERF domain-containing transcription factor [Populustrichocarpa] |
| Unigene0006534 | 3.1 | -2.0 | Basic helix-loop-helix DNA-binding family protein [Theobroma cacao] |
| Unigene0006504 | 3.9 | -3.6 | transcription factor bHLH96-like [Vitisvinifera] |
| Unigene0023715 | 7.8 | -2.8 | transcription factor bHLH74 [Vitisvinifera] |
| Unigene0009434 | 2.9 | -2.2 | BES1/BZR1-like protein [Medicagotruncatula] |
| Unigene0007439 | 5.7 | -4.3 | Trihelix transcription factor GT-3b-like [Cicer<br>arietinum] |

## Acknowledgements

This work was supported by the Project of Science and Technology in Yunnan: model construction and demonstration of wildlife tending for anticancer material *M. tenacissima* (grant No. 2016RA009), the Project of Young and Middle-aged Talent of Yunnan Province (grant No. 2016PY068), and the National Natural Science Foundation of China (grant No. 81660636 and 81660635).

## Supplemental Data

**Supplementary table 1 Differentially expressed genes that were up-regulated in the CK vs. T1comparison and down-regulated in the T1 vs. T2 comparison**

**Supplementary table 2 Differentially expressed genes that were down-regulated in the CK vs. T1comparison and up-regulated in the T1 vs. T2 comparison**

**Supplementary table 3 Significantly differentially expressed genes identified in the CK vs. T1 comparison that were annotated to KEGG pathways**

**Supplementary table 4 Significantly differentially expressed genes identified in the CK vs. T2 comparison that were annotated to KEGG pathways**

**Supplementary table 5 Significantly differentially expressed genes identified in the T1 vs. T2 comparison that were annotated to KEGG pathways**

**Supplementary table 6 Differentially expressed transcription factors identified in the CK vs. T1 comparison**

**Supplementary table 7 Differentially expressed transcription factors identified in the CK vs. T2 comparison**

**Supplementary table 8 Differentially expressed transcription factors identified in the T1 vs. T2 comparison**

**Supplementary table 9 Genes that were up-regulated under drought stress and down-regulated after re-watering treatment**

**Supplementary table 10 Log2 fold change data for the RNA-seq and qRT-PCR analyses.**

